# High MHC gene copy number maintains diversity despite homozygosity in a Critically Endangered single-island endemic bird, but no evidence of MHC-based mate choice

**DOI:** 10.1101/2020.02.03.932590

**Authors:** Martin Stervander, Elisa G. Dierickx, Jack Thorley, M. de L. Brooke, Helena Westerdahl

**Author notes:** **Corresponding authors:** Martin Stervander, Helena Westerdahl. These authors contributed equally (Martin Stervander, Elisa G. Dierickx, Jack Thorley).

## Abstract

Small population sizes can, over time, put species at risk due to the loss of genetic variation and the deleterious effects of inbreeding. Losing diversity in the major histocompatibility complex (MHC) could be particularly harmful, given its key role in the immune system. Here, we assess MHC class I (MHC-I) diversity and its effects on mate choice and survival in the Critically Endangered Raso lark *Alauda razae*, a species restricted to the 7 km^2^ islet of Raso (Cape Verde) since ~1460, whose population size has dropped as low as 20 pairs. Exhaustively genotyping 122 individuals, we find no effect of MHC-I genotype/diversity on mate choice or survival. However, we demonstrate that MHC-I diversity has been maintained through extreme bottlenecks by retention of a high number of gene copies (at least 14), aided by co-segregation of multiple haplotypes comprising 2–8 linked MHC-I loci. Within-locus homozygosity is high, contributing to comparably low population-wide diversity. Conversely, each individual had comparably many alleles, 6–16 (average 11), and the large and divergent haplotypes occur at high frequency in the population, resulting in high within-individual MHC-I diversity. This functional immune gene diversity will be of critical importance for this highly threatened species’ adaptive potential.

## 1 Introduction

Islands have intrinsic geographic characteristics (isolation, small area) that underlie the exacerbated genetic threats to insular species. Isolation can make gene exchange difficult, which, coupled to the often small size of island populations, can increase these risks (Ferreira et al., 2016; Richard Frankham, 1998). One striking example of the genetic threats to insular species is that of the last woolly mammoth *Mammuthus primigenius* population, where 300 individuals surviving on Wrangel Island are thought to have ultimately disappeared because of an accumulation of deleterious mutations that led to “genomic meltdown” (Rogers & Slatkin, 2017) (even flying animals like birds and bats can exhibit strong philopatry and limited dispersal abilities (e.g. Chaverri & Kunz, 2011; R. Frankham, 1997)_)_. Indeed, the maintenance of genetic diversity is crucial for fitness and survival, at the individual, population and species level (Richard Frankham, Ballou, & Briscoe, 2002). Genetic diversity increases the viability of recently translocated populations and helps species respond to environmental changes (Hughes, Inouye, Johnson, Underwood, & Vellend, 2008; Johnson et al., 2010; Keller & Waller, 2002; Nair, 2014; Reusch, Ehlers, Haemmerli, & Worm, 2005; Romiguier et al., 2014; Tollington et al., 2013; Wright, Gillman, Ross, & Keeling, 2009). The maintenance of genetic diversity in an island population is therefore particularly important. There is evidence that species on smaller landmasses (islands) have lower rates of molecular evolution than species on larger landmasses (Tollington et al., 2013; Wright et al., 2009). This suggests that confining species to small refugia reduces the rate of microevolution, which could limit the species’ ability to adapt to environmental changes (Tollington et al., 2013; Wright et al., 2009).

It is on one of these small island refugia, the uninhabited 7 km^2^ islet of Raso (16° N 24° W), that the Critically Endangered Raso lark *Alauda razae* survives. There the population was counted intermittently during the 20^th^ century, and fluctuated, rising after higher rainfall (Donald, De Ponte, Pitta Groz, & Taylor, 2003). Since 2001, the population has been monitored annually in October–December. During this period, the lark has shown dramatic variation in population size, from a low of about 57 individuals in 2004, to a high of 1,500–1,550 individuals in 2011, 2012 and 2017 (Brooke, 2019). The Raso lark is a sexually dimorphic species (the males are larger than the females) that breeds irregularly, following the similarly irregular rainfalls (Donald & de L. Brooke, 2006). When breeding is possible, most individuals live and forage in socially monogamous pairs on the plains of Raso (Donald & de L. Brooke, 2006). Only once, among several hundred breeding pairs observed, have we seen a breeding male consorting with two females. When not breeding, the birds are mainly found foraging in large flocks both on the plains and the plateaus of the island.

Potentially detrimental to the Raso lark’s persistence are the population bottlenecks through which the species has passed. A species with a small effective population size is subject to three types of genetic risk. The first is inbreeding depression, which is reduced fitness due to the increase in homozygotes caused by mating between relatives (Edmands, 2007; O’Grady et al., 2006). Dierickx, Sin, et al. (2019) found three closely related (sibling or parent–offspring) pairs of Raso larks out of 26 sampled, suggesting that the species might indeed be at risk of inbreeding. The second type of genetic risk is the loss of potentially adaptive genetic variation which limits the species’ ability to adapt in response to environmental changes (O’Grady et al., 2006; Reed, 2005; Tollington et al., 2013; Wright et al., 2009) such as, for example, climate change, to which Cape Verde – as an arid country in the Sahel region and a small island state – is doubly at risk (Dierickx, Robinson, & Brooke, 2019; Ministry of Environment Housing and Territory Planning of Cape Verde, 2011). The third is deleterious allele accumulation, also called “mutational meltdown”, due to the fact that selection is weaker in small populations than in large populations (O’Grady et al., 2006; Reed, 2005).

In the context of a Critically Endangered species potentially exposed to genetic risk, characterizing the species’ Major Histocompatibility Complex (MHC) is of theoretical interest, as well as of use for conservation. Indeed, the MHC is a very diverse part of the genome that typically contains multiple gene copies, and therefore is likely to maintain at least some genetic diversity, even if variation at the rest of the genome is heavily depleted. Furthermore, an organism’s immune system is crucial to its fitness and hence to survival. One final reason is our interest in investigating MHC-based mate choice in this species. We hypothesize that, given the high risk of inbreeding that Raso larks face (Dierickx, Sin, et al., 2019), individuals are under strong selective pressure to develop a mating strategy that maximizes the genetic diversity of their future offspring. Basing mate choice on MHC genotype could be advantageous for the two reasons mentioned above: it might be one of the few indicators of genetic diversity left in the genome, and it plays a key role in the immune system.

In the present study we characterize MHC class I (MHC-I) in Raso larks and measure MHC-I genetic diversity. We then test if there is an assortative MHC-I based mate choice and investigate whether MHC-I genetic diversity is associated with survival.

## 2 Material and Methods

### 2.1 Sample collection and blood sampling

Each year since 2004 a two-person team has spent two to three weeks on Raso in November or early December, catching and ringing new flying birds, and also ringing nestlings and juveniles (<3 months old), the latter recognized by their browner plumage with broader pale feather edgings. Captured birds, readily sexed thanks to sexual size dimorphism (Donald et al., 2003), received an individually-numbered metal ring and a unique combination of three Darvic colour rings which allow individual identification in later years (assuming the bird survives). In addition, at the time of first capture, a blood sample was taken by pricking the brachial vein and collected onto EDTA-moistened filter paper. After air drying, the blood-stained paper was stored at ambient temperature in the field and then at –80°C after return to the UK. The blood samples were obtained by a licensed bird ringer (M. de L. Brooks: British Trust for Ornithology permit A 1871 MP) without damaging the health of the birds and with permission from the Cape Verdean authorities (Direcção Nacional do Ambiente).

The proportion of the population that was ringed varied from roughly 30–60 percent, tending to be lower when the population was higher. Two colour-ringed birds were considered to be breeding together when they jointly attended a nest with eggs and/or young, or when they were both seen feeding a recently-fledged juvenile. A male and female socially consorting but without evidence of breeding together are not considered paired for the purposes of the present paper.

### 2.2 Primer design and initial evaluation

We initially designed new primers to amplify exon three of MHC class I, based on aligning available passerine MHC class I and genomic sequences. We aimed at anchoring primers in relatively conserved regions and added degenerated bases to account for sequence variation. We furthermore attempted to amplify as much of the exon as possible by anchoring the primers in flanking intron sequence. We evaluated three new forward and four new reverse primers in twelve possible primer combinations, as well as the primer combination HNalla (O’Connor, Strandh, Hasselquist, Nilsson, & Westerdahl, 2016) and HN46 (Westerdahl, Wittzell, von Schantz, & Bensch, 2004). For primer sequences, see Table S1.

The initial evaluation was performed in 10 μl PCR reactions on samples of great reed warbler, Raso lark, blue tit *Cyanistes caeruleus*, and zebra finch *Taeniopygia guttata*, using the Qiagen Multiplex PCR Kit (Qiagen Inc., Hilden, Germany). We included 5 μl Qiagen Multiplex PCR Master Mix, 0.2 μl each of 10 μM forward and reverse primer, 2 μl template DNA (5–10 ng/μl) and 2.6 μl water. The PCRs were run with an activation at 95°C for 15 min; 35 three step cycles with denaturation at 94°C for 30 s, annealing at varying temperatures for 90 s (Table S2), and extension at 72°C for 90 s; with a final extension at 72°C for 10 min. The PCR products were the run on a 2% agarose gel stained with GelRed (Biotium, Fremont, CA, USA), and visually inspected in UV lighting (Table S2, Figure S1).

Based on the above, three combinations of the new primers (3F_us30-us4 + 3R_ds24-ex2, 3F_us35-us8 + 3R_ds20-ex6, and 3F_us35-us8 + 3R_ds24-ex2) and HNalla + HN46 were selected for further evaluation by producing sequences for six Raso larks in a shared amplicon library sequenced on an Illumina Miseq, after which the results were evaluated (see below) and a final primer combination was selected for creating a new amplicon library in which 130 samples were included.

### 2.3 Library preparation

We prepared two amplicon libraries for Illumina sequencing using a two-step amplification. First, six (first library) and 130 (second library) individual samples were amplified using four primer pairs (first library; see above) and 3F_us30-us4 + 3R_ds24-ex2 (second library) modified with 5’-overhangs designed to match the Illumina sequencing adapters and molecular identifiers (MIDs) of the Nextera® XT v2 Index Kit (Illumina Inc., San Diego, CA, USA). The second library included eight technical duplicates, four of which were independent extractions from different blood samples from different trapping occasions of the same individual, and four of which the same sample was used twice. The reactions comprised 25 μl and used 25 ng template DNA, 0.5 μM of each primer, and 12.5 μl 2X Phusion High-Fidelity PCR Master Mix (ThermoFisher Scientific, Waltham, USA). The PCR was initiated with a 30 s denaturation step at 98°C followed by 25 cycles of 10 s denaturation at 98°C, 10 s annealing at 66.8°C, and 15 s elongation at 72°C. A 10 min final extension 72°C completed the program.

The PCR product was cleaned with Agencourt AMPure XP-PCR Purification Kit (Beckman Coulter, Indianapolis, USA), following the manufacturer’s instruction with some modifications: The ratio of PCR product to beads was 1:0.8, 80% ethanol was used in the bead cleaning steps, and the elution was made with 43 μl double-distilled water, which incubated at room temperature for two minutes. An aliquot of the clean PCR product was run on a 2% agarose gel, to verify fragment length and to roughly estimate concentration of the PCR product based on band intensity. The individual PCR products were then differentially evaporated at room temperature, to achieve even concentrations.

In the second PCR, MIDs were added to each of the samples, as well as 360 (first library) and 254 (second library) unrelated samples prepared with various primer combinations. We used unique combinations of forward and reverse Illumina indices for each sample, using the Nextera XT v2 Index Kit (Illumina Inc., San Diego, CA, USA). PCRs were run in 50 μl reactions and contained 25 μl 2X Phusion High-Fidelity PCR Master Mix (ThermoFisher Scientific, Waltham, USA), 5 μl of each index primer, and—depending on estimated concentration—5, 7.5, 10, or 15 μl cleaned PCR product. The PCR was initiated with a 30 s denaturation step at 98°C followed by eight cycles of 10 s denaturation at 98°C, 15 s annealing at 62°C, and 15 s elongation at 72°C. A 10 min final extension at 72°C completed the program.

The indexed amplicons were cleaned with Agencourt AMPure XP-PCR Purification Kit (Beckman Coulter, Indianapolis, USA) as specified above, but with a ratio of PCR product to beads at 1:1.12. The cleaned PCR products were checked on a 2% agarose gel, and quantified using a Quant-iT PicoGreen dsDNA Assay Kit (ThermoFisher Scientific/Invitrogen, Waltham, USA) modified for a 96-well plate, measured on a plate reader.

We pooled an equimolar quantity of each of 384 samples into four pools (for the first library, including the four primer combinations and six individuals) or six pools (for the second library, including the final sequencing), depending on amplicon length, concentration, and primer combination. These pools were then quantified with Qubit Broad Range and High Sensitivity kits (ThermoFisher Scientific, Waltham, USA), after which we ran them on a Bioanalyzer DNA 2100 chip (Agilent, Santa Clara, CA, USA) for validation of quality and size. In a final step, equimolar quantities of all pools were combined in a 20 nM library, which was sequenced with 300 bp paired-end Illumina MiSeq sequencing (Illumina Inc., San Diego, CA, USA) at the DNA sequencing facility of the Department of Biology, Lund University.

### 2.4 Bioinformatic pipeline

The sequence output of each sample was first trimmed using Cutadapt (Martin, 2011), based on base quality of the 3’-end (−q 15) followed by linked adapter validation and removal of the paired reads (−a, −A). Trimmed paired reads were then imported to DADA2 v. 1.0.3 (Callahan et al., 2016) in R v 3.5.0 (R Core Team, 2018) where we filtered with fastqPairedFilter, adjusting the criterion for expected error rate for reverse reads from 2 to 5 (maxEE=c(2,5)); dereplicated with derepFastq; learned error rates with the learnErrors function called through dada; and merged read pairs with mergePairs. Thereafter, a sequence table was created with makeSequenceTable, after which sequence length distributions were checked and any spurious sequences of very deviating lengths were excluded, and the remaining sequences were filtered for sequence chimeras using removeBimeraDenovo.

### 2.5 Evaluation and selection of primers

The output from dada2 based on the six individuals amplified with four primer pairs in the first library was scrutinized and cross referenced. HNalla + HN46 rendered significantly fewer alleles, whereas most alleles were recovered by the three other primer combinations, some with drastically varying coverage, however. For library 2, we selected the primer pair 3Fus35us8 + 3Rds20ex6, which generated >500 reads in ≥1 of 6 test samples for alleles A–V (see Figure 1, Table S3) with the exceptions of alleles L, S, and T (all of which did not occur in those samples). Further, additional allele 1 occurred with 179–240 reads and additional allele 2 with only 65–102 reads (see Figure S2), whereas they (and allele V) occurred in high frequency (>1000 reads) with two other primer combinations (which, in turn, did not amplify other alleles well). Finally, another seven alleles (additional putative alleles 3–9; Figure S2) were amplified at very low frequency.

**Figure 1.**
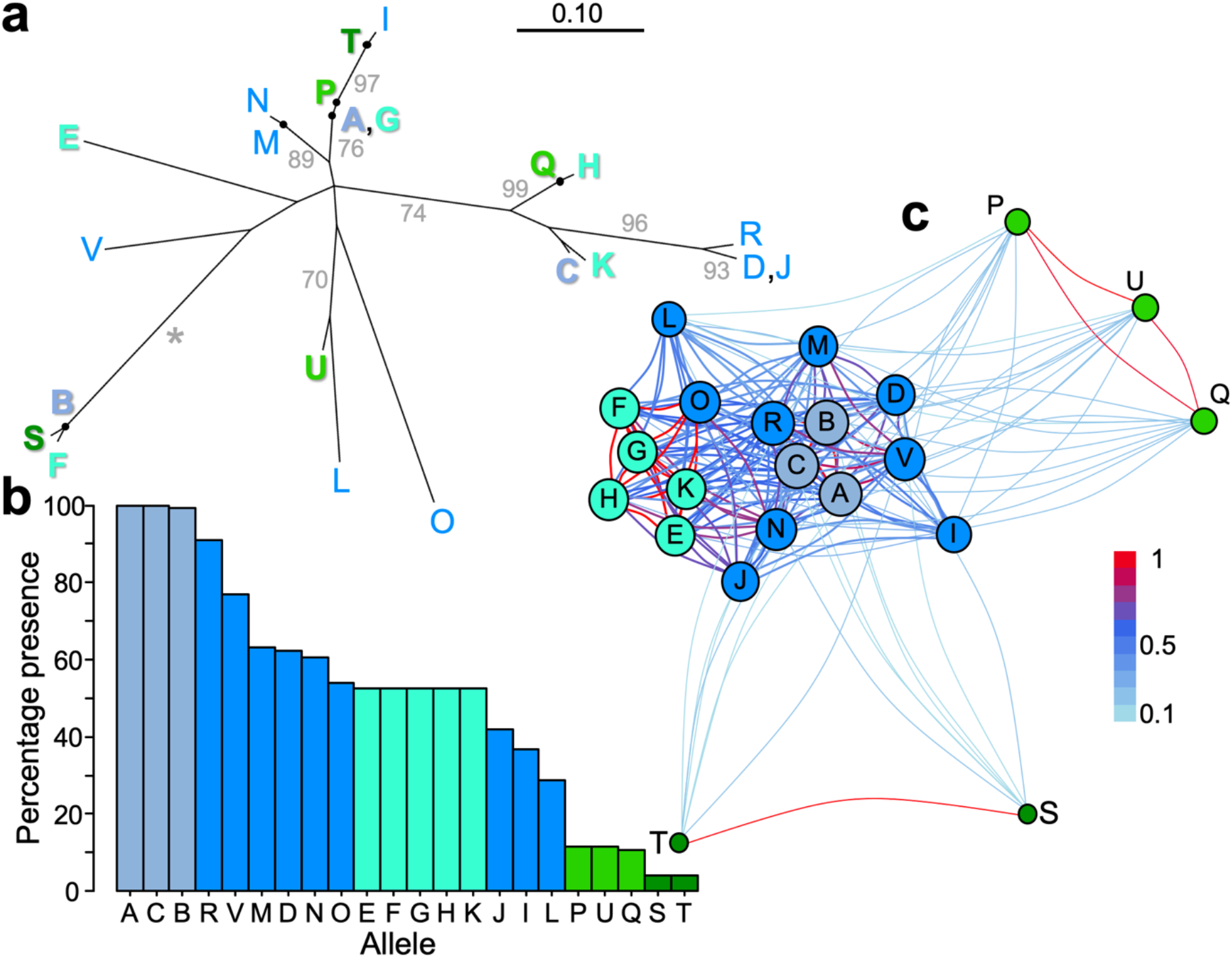
MHC-I allele characteristics in the Raso lark *Alauda razae*, based on 122 genotyped individuals. Colours other than medium blue indicate fixed alleles (*ABC* in grey-blue) or co-segregating allele blocks (*EFGHK*, *PQU*, and *ST* in different shades of green) and correspond to Table S3 and Figure S4. (**a**) Unrooted MHC-I amino acid allele tree, computed with the LG substitution model. Allele names are placed at tips or internal terminal nodes (marked with black circles). Two pairs (AG and DJ) do not differ by any non-synonymous mutations (see Figure S4). Fixed or almost fixed alleles, as well as alleles in co-segregating clusters, labelled in bold. Bootstrap values >70%, based on 1,000 replicates, are written with grey font (at the branch upstream of supported node), with 100% indicated by an asterisk. See Figure S2 for a corresponding tree, including an additional nine alleles that are certainly (1–2) or likely (3–9) functional, but yielded too few reads with our primer pairs to enable confident allele calls (see Material and Methods). (**b**) Allele frequency (% of 122 birds) in the populations. (**c**) Network graph illustrating the allele co-occurence. Nodes display each allele, sized according to its frequency in the sampled population. Edges indicate alleles that are present together within individuals, coloured according to the proportion of allele sharing; for example, S and T are present in relatively few individuals, but when present they co-occur within individuals at high rates (> 0.9).

### 2.6 Further filtering and allele calls

The combined output of library 1+2 from dada2 comprised 136 samples with an effective coverage (number of reads used by dada2) between 0 (one failed sample) and 119,942, with 50% of samples having 20,044–26,070 (median 22,322) reads. It was further filtered as follows: (1) Five samples with less than 5,000 reads were excluded. (2) Alleles for which the exon contained an uninterrupted open reading frame were accepted if the average read count for individuals in which they were called exceeded 500. In addition, there were 11 alleles with very low read counts, which were not included since the call whether present in an individual would not be reliable. We made an exception for two of those 11 low-frequency alleles (R, V), because (a) they had considerably higher read counts in the six samples from the first library, using the final primer pair; (b) these two alleles were called consistently between multiple primer pairs in library 1, with some primer pairs producing high read counts; and (c) there was concordance between technical duplicates, with the exception of a single allele call in one duplicate with few total reads. (3) For an allele to be called in an individual, its effective coverage had to meet either (a) a threshold value of 10% of the average coverage for that allele for samples in which it was present, or (b) a threshold value of within-individual read frequency of 5% (i.e. >5% of all reads in that individual had to belong to that allele).

Allele calls were compared between the eight pairs of technical replicates, and repeatability calculated as 2 × shared allele calls / (allele calls of replicate A + allele calls of replicate B). Among the remaining 122 unique samples used for analyses, there was no effect of coverage on number of alleles (F_1,120_ = 0.010, r^2^ = 0, p = 0.76; Figure S3). We lacked individual data (sex, age, morphometric) for eight samples, which were included in overall diversity analyses, but excluded from analyses of survival and mate choice.

### 2.7 MHC-I diversity and test of trends over time or sex effects

MHC-I diversity was computed with PhyML (Guindon & Gascuel, 2003) as the total length of a phylogenetic tree of the amino acid sequences of an individual’s alleles, using the LG substitution model (Le & Gascuel, 2008). The relationship between number of alleles and MHC-I diversity was explored by linear models.

Because few individuals were ringed as nestlings or recently-fledged juveniles, and because Raso larks cannot be aged accurately from plumage after their complete post-juvenile moult around three months of age, we instead approximated the age of individuals when first captured using information on claw damage. Previous data from the population indicates that, while birds less than two years old have undamaged feet, approximately one third of all birds known to be at least two years of age show clear signs of toe or claw damage. We therefore assumed that birds with claw damage on their first capture were two years old, whereas any birds with no toe damage were in their first year of life (Age 1). This is most likely to be strictly true after a year of strong population growth (e.g. 125% increase from 2009 to 2010). The data on the 114 individuals is compiled in Supporting Information 2.

Temporal trends were tested with a linear regression (MHC-I diversity ~ inferred birth year) and differences between males and females with a two-tailed t-test.

### 2.8 Randomisation tests of (dis-)assortative mating

We used randomisation tests to determine whether Raso larks mate (dis-)assortatively. Of the 114 birds with individual data that were genotyped, there were 46 pairings where both the male and female bird in a pair were genotyped. For each pairing (n = 46), we calculated the proportion of shared MHC-I alleles and the pairwise mean amino acid distance. The preliminary analyses identified two alleles that were fixed in the population, another allele present in all but one sample, and three co-segregating blocks of alleles (see Results; Figure 1b-c). Because of the potential influence of these co-segregating alleles on estimates of allele sharing we calculated shared allelic diversity in two ways; treating the co-segregating blocks as separate alleles (termed ‘allele sharing α’), and as single allelic blocks (‘allele sharing β’). The empirical values for each of the metrics were then compared to frequency distributions of the mean values generated from 9,999 permutations of 46 randomly selected parent pairs (including the observed value generates a distribution of 10,000 values). For each randomised mating within a permutation, we assumed that females could mate with any genotyped male present in the population in the year of mating, sampling with replacement. We set the year of mating as the first year in which a pair were observed together, and, because Raso larks are socially monogamous and pairs frequently mate together in successive years, we did not allow females to mate with males that were present in the population but which were known to have been paired to a female with whom they had mated previously (i.e. removing the confounding effect of breeding history; see Table S4 for breakdown of birds in each year). Observed values falling outside the 2.5–97.5% confidence intervals of the frequency distribution for each metric would indicate significant departures from random mating at an alpha level of 0.05.

### 2.9 Survivorship

We used a Cox proportional hazards analysis to investigate the effect of MHC-1 diversity on age-related survivorship, where MHC-1 diversity represents the total tree length per genotype. Age categorisation was carried out using information on claw damage and plumage (juveniles) as described above. The Cox analysis fitted the time at death as the response term, with the age category, sex, and MHC-1 diversity specified as fixed effects. Birds that were last seen in 2017 and 2018 were right censored to account for uncertainty in re-sighting, though annual re-sighting rate is high for this population (c. 88%; own data). All statistical tests were performed in R v. 3.6.1 (R Core Team, 2019).

## 3 Results

### 3.1 MHC-I diversity

From a total of 130 samples, including eight technical replicates, we identified 22 MHC-I alleles (Figure 1; Table S3; Supporting Information 3), many of which were present at high levels in the sampled population (Figure 1b), and several of which co-occurred within individuals (Figure 1c). Of the 22 alleles, which we named alphabetically following decreasing across-population allele read depth in the second library, 20 (alleles A–Q, S–U) occur with higher within-individual (or within-sample) frequency (read depth) whereas two (alleles R, V) occur with markedly lower within-individual frequency (Table S3). We consider these two alleles valid, but under-amplified by our primers, which means that we risk false negatives, when fewer than sufficient reads qualify for allele calls in certain individuals (see Methods). The repeatability among 75 pairwise allele comparisons in eight replicated samples was high, 99%: there was a single difference when the low within-individual frequency allele R was not called in a replicate that had the sixth lowest coverage (13,105×) out of all 130 samples.

Among the 122 individuals, the number of different MHC-I alleles per individual varied between six and 16 alleles, with an average of 11.2 alleles (excluding *RV* alleles: 4–15, average 9.6). Three alleles (A, B, C) occurred in almost all individuals (100%, 99%, 100%), and a co-segregating block of five alleles (E–H and K) occurred in 52% of the individuals (Figure 1b-c). To evaluate the genomic architecture of the two common co-segregating allele blocks *ABC* and *EFGHK*, we calculated the ratio of the average number of sequence reads for *EFGHK* alleles over the average number of sequence reads for *ABC* alleles in individuals possessing both blocks, and found a clearly bimodal distribution with averages 0.52 (standard deviation = 0.06, n = 51) and 1.06 (standard deviation = 0.06, n = 13; Figure 2a), indicative of copy number variation (one or two sets) of the *EFGHK* alleles, consistent with two main haplotypes comprising *ABC* and *ABCEFGHK* and three genotypes: homozygous for *ABC*, heterozygous for *ABCEFGHK*|*ABC*, and homozygous for *ABCEFGHK*.

**Figure 2.**
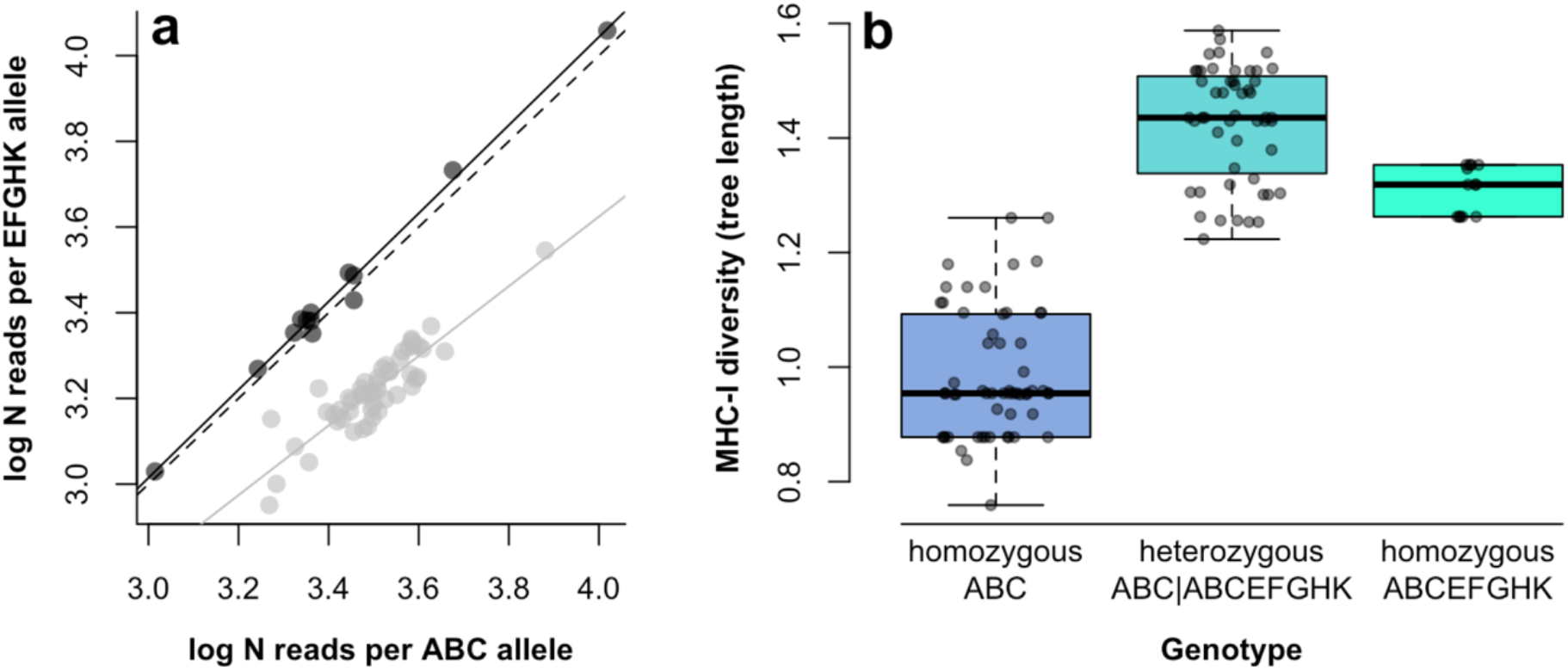
(**a**) Relationship between allele read depth of *ABC* alleles (log number of reads, x-axis) and *EFGHK* alleles (log number of reads, y-axis) in individuals with *EFGHK* alleles. The individuals are grouped according to a bimodal distribution of the ratio of average allele read depth for *EFGHK* alleles over *ABC* alleles, with individuals interpreted as heterozygous for *ABCEFGHK*|*ABC* coloured grey (ratio 0.44–0.76, average 0.52, standard deviation 0.06; all but two individuals ≤0.60). and individuals interpreted as homozygous for *ABCEFGHK* coloured black (ratio 0.94–1.14, average 1.06, standard deviation 0.06; all but two individuals ≥1.0). Solid lines correspond to linear regression slopes within each genotype, dashed line indicates a 1:1 ratio. (**b**) Distributions of MHC-I diversity (measured as total allele tree length) depending on genotype for the *ABCEFGHK*|*ABC* loci. All three genotypes significantly differ in MHC-I diversity, with heterozygotes on average most diverse.

We also found another co-segregating block composed of two rarer alleles (S, T), occurring in five individuals (Figure 1b-c; one individual run as a technical replicate). Finally, there was an intermediately rare block of alleles (P, Q, U) that co-segregated in 11 out of 12 individuals (Q not detected in one of the samples; Figure 1b-c).

Out of the 11 alleles or co-segregating blocks other than *ABC* and *EFGHK*, six deviated significantly from expected random association to *ABCEFGHK*|*ABC* genotypes (Table S5). Five alleles/blocks (three significantly so) associated with the *ABC* haplotype and were never observed in *ABCEFGHK* homozygotes, whereas two (both significantly) associated with the *ABCEFGHK* haplotype and was under-represented in *ABC* homozygotes (Table S5). Allele O co-segregated with *EFGHK* in all 64 *ABCEFGHK*|*ABC* heterozygotes and *ABCEFGHK* homozygotes, but was also observed in 2 of 58 *ABC* homozygotes (Table S5).

The raw distances between alleles ranged 1–42 nucleotides or 0–26 amino acids (Figure S4). However, since the differences between biochemical properties of amino acids vary greatly in magnitude, we defined amino acid *divergence* as the patristic distances (branch length distance) computed with the LG substitution model (Le & Gascuel, 2008): the average divergence between all alleles was 0.337 ± 0.009. The fixed (or almost fixed) A, B and C alleles were found in different parts of the allele tree (Figure 1a), with an average divergence of 0.364 ± 0.068. This was true also for the co-segregating blocks *EFGHK* (0.356 ± 0.039), *PQU* (0.257 ± 0.034), and *ST* (0.420).

MHC-I *diversity* in an individual is determined by the number of alleles and their degree of dissimilarity, a metric that we defined as the total branch length of a phylogenetic tree comprising an individual’s alleles (computed with the LG substitution model (Le & Gascuel, 2008)). Using this definition, the average within-individual MHC-I diversity was 1.205 ± 0.021, but number of alleles was a good proxy for diversity, explaining 90% of the variation (adjusted r^2^) in a simple linear model (diversity ~ N alleles; b = 0.075, F_1,120_ = 1,086, p < 2.2×10^−16^). Individuals with the five co-segregating *EFGHK* alleles had on average 14.2 alleles ± standard error (SE) 0.8, which is 5.9 alleles (71%) more than individuals without the five *EFGHK* alleles (average 8.3 ± 0.2 alleles). When separating *EFGHK* bearers into homozygotes for *ABCEFGHK* and heterozygotes for *ABCEFGHK*|*ABC*, heterozygotes had higher diversity than *ABCEFGHK* homozygotes, both of which had higher diversity than *ABC* homozygotes (diversity ~ N alleles + *ABCEFGHK*|*ABC* genotype; F_3,118_ = 373.1, p < 2.2×10^−16^; Figure 2b). There was no effect of sex (t_1,112_ = 0.38, r^2^ = 0, p = 0.70) on MHC-I diversity, neither was there any temporal trend based on inferred year of birth (t_1,112_ = –0.89, r^2^ = 0, p = 0.37; Figure S5).

### 3.2 No MHC-I based mate choice

There was no difference in the total allele number of males and females, irrespective of whether the *RV* alleles were included (male mean = 11.30, SD = 3.06; female mean = 10.95, SD = 2.90; Welch’s t-test t_109.2_ = –0.62, p = 0.54), or excluded (male mean = 9.67, SD = 3.20; female mean = 9.25, SD = 3.04; Welch’s t-test t_109.5_ = –0.71, p = 0.48).

Raso lark pairs typically shared a high proportion of alleles (mean = 0.67, SD = 0.2), but this was not any more or less than that expected under random mating, irrespective of how we handled the co-segregating blocks (Table 1; Figure 3a-b), or treated the *RV* alleles (Table S6). Randomisation tests likewise found no evidence for assortative or disassortative mating according to divergence (mean amino acid distance; Table 1; Figure 3c; Table S6). Though our sample size was modest, in all cases, the empirical mean of the different complementarity metrics sat far from the 2.5% tails of the distribution of means generated under random mating (0.23 < p < 0.79).

**Table 1.**
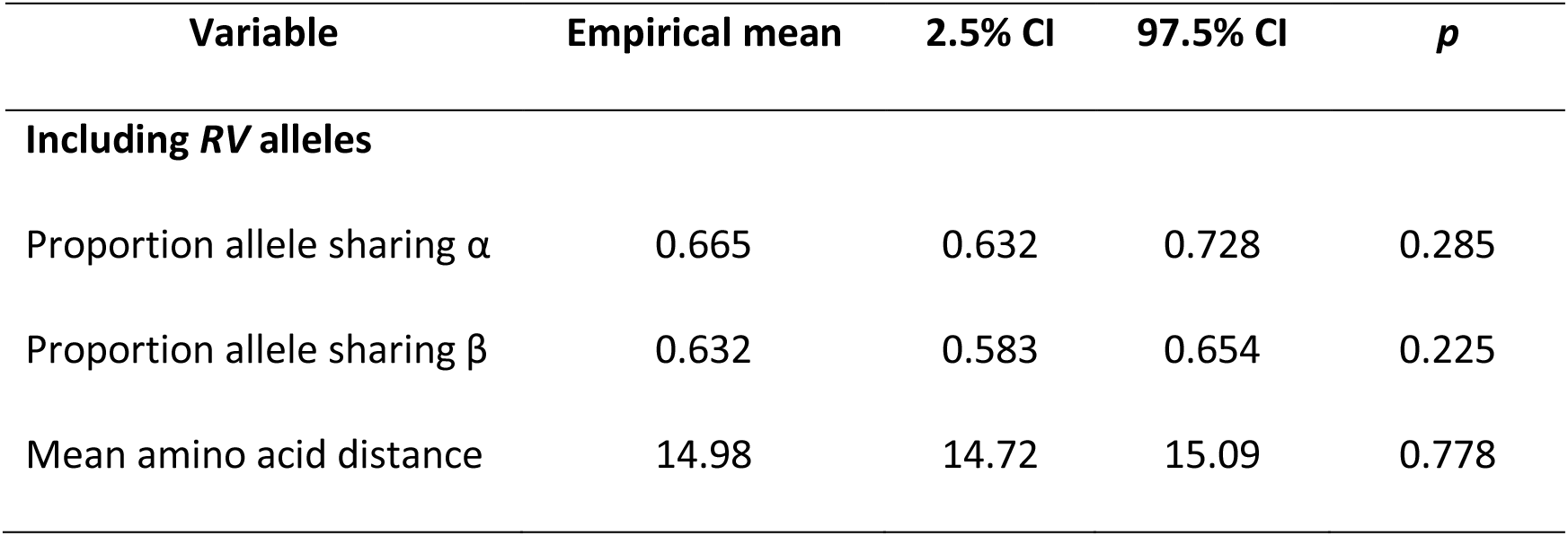
Randomisation testing of (dis-)assortative mating in Raso larks based on allelic variation in MHC-I. The empirical means calculated from observed pairings were compared to randomly assigned pairings generated from 9,999 permutations; significant departures from random mating would fall outside the 95% confidence intervals (CI). α and β refer to the whether the co-segregating block of five alleles were treated as five separate alleles (α), or a single allelic block (β). *p* is the ranked position of the observed mean in the distribution. Note that here, all results to randomisations that included the *RV* alleles; for results excluding *RV* alleles, see Table S3.

| Variable | Empirical mean | 2.5% CI | 97.5% CI | $p$ |
| --- | --- | --- | --- | --- |
| **Including *RV* alleles** |  |  |  |  |
| Proportion allele sharing $\alpha$ | 0.665 | 0.632 | 0.728 | 0.285 |
| Proportion allele sharing $\beta$ | 0.632 | 0.583 | 0.654 | 0.225 |
| Mean amino acid distance | 14.98 | 14.72 | 15.09 | 0.778 |

**Figure 3.**
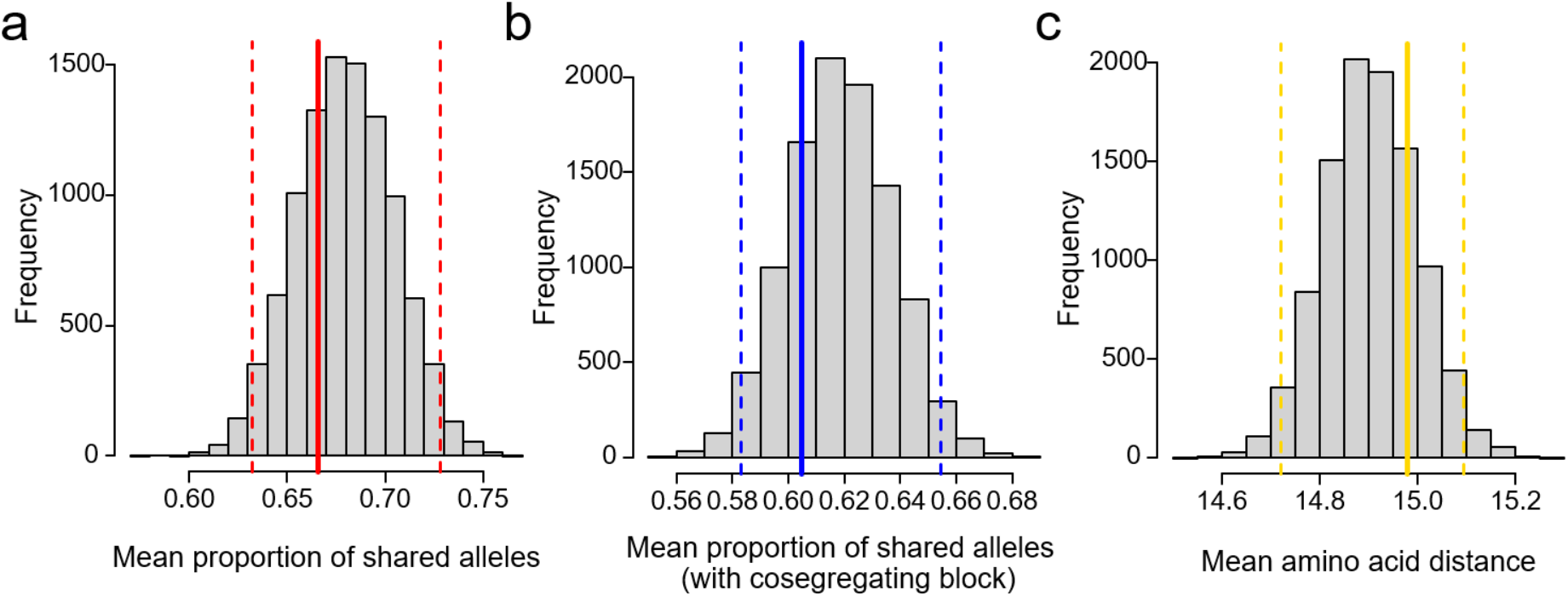
Randomisation testing of (dis-)assortative mating based on allelic variation. (**a**) Mean proportion of shared alleles. (**b**) Mean proportion of shared alleles, treating co-segregating blocks as single units. (**c**) Mean amino acid distance. Solid lines show empirical means from observed pairings. Frequency distributions show mean values generated from 9,999 permutations of random pairing (including the empirical data = 10,000). The two-tailed 95% confidence intervals (dashed lines) display cut-offs for significant departures from random mating. N = 46 pairings. For corresponding analyses excluding the *RV* alleles, see Figure S7.

### 3.3 No MHC-I based fitness effects

Raso larks display high survivorship (Figure S6; Dierickx, Robinson, et al. (2019)). Nevertheless, the categorisation of individuals into general age groupings on the basis of claw damage and information on whether the individual was ringed as a *pullus* (in the nest) provided support for a graded effect of age on survival (χ^2^_2_= 7.54, p = 0.029; Table 2), where birds ringed as a *pullus* survived longer than adult birds without claw damage on first capture, which in turn tended to survive longer than adult birds with claw damage on first capture (Figure S6). After accounting for this age pattern, we found no effect of the MHC-I diversity on survival in either sex (Table 2).

**Table 2.**
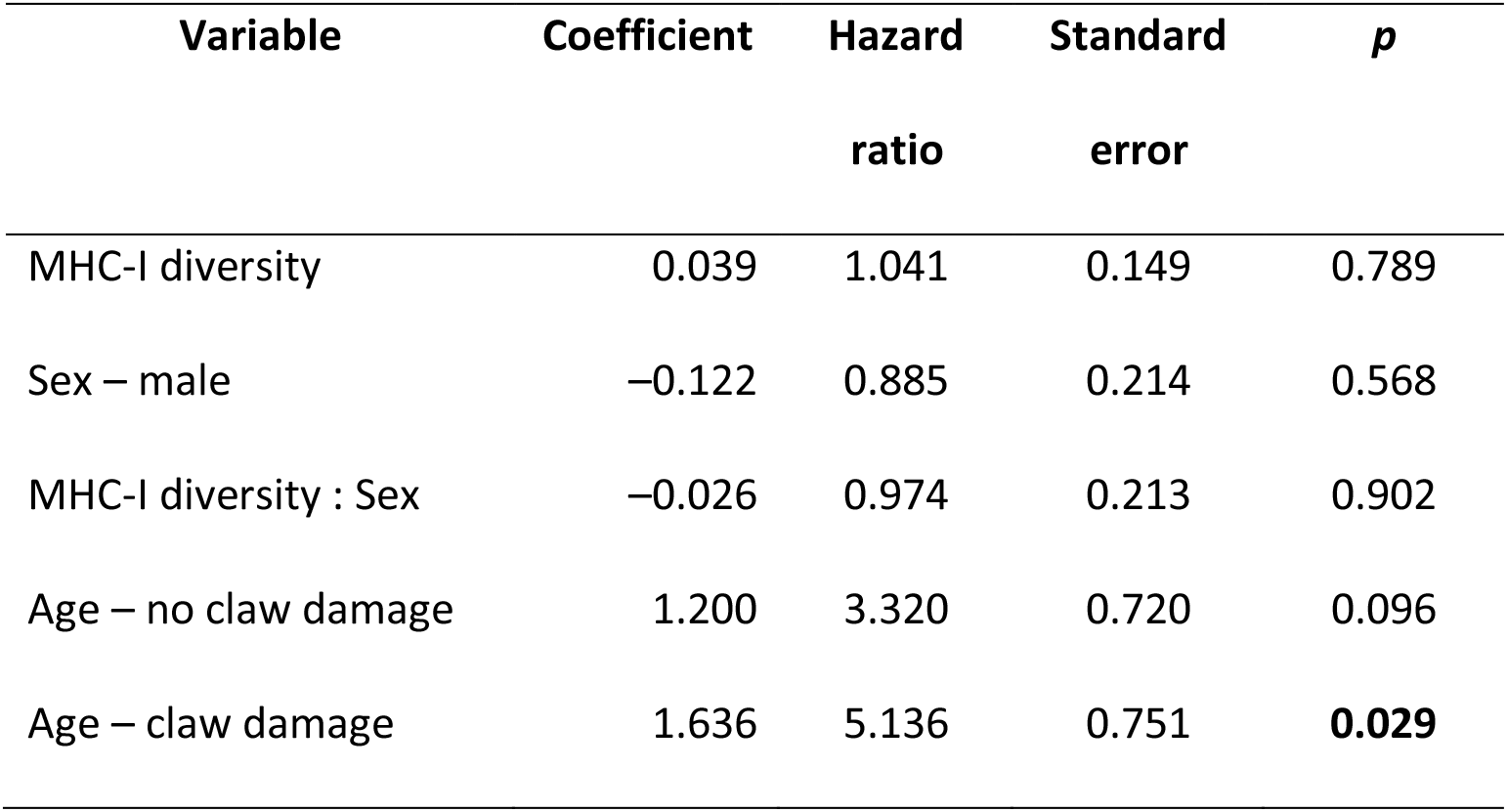
Survivorship of Raso larks according to MHC-I diversity. A hazard ratio of 1.04 implies that 1 standardised unit increase in MHC-I diversity, measured as total tree length of a genotype, increases the risk of hazard by 4%, but note that only age-related variables were significant or near-significant. Age category refers to a general age categorisation according to claw damage (older birds are more likely to have claw damage) and information on whether a bird was ringed as a *pullus* (in the nest). Reference levels for the sex and age category covariates are ‘female’ and ‘ringed as a *pullus*’, respectively.

| Variable | Coefficient | Hazard<br>ratio | Standard<br>error | *p* |
| --- | --- | --- | --- | --- |
| MHC-I diversity | 0.039 | 1.041 | 0.149 | 0.789 |
| Sex – male | −0.122 | 0.885 | 0.214 | 0.568 |
| MHC-I diversity : Sex | −0.026 | 0.974 | 0.213 | 0.902 |
| Age – no claw damage | 1.200 | 3.320 | 0.720 | 0.096 |
| Age – claw damage | 1.636 | 5.136 | 0.751 | **0.029** |

## 4 Discussion

Raso larks have low population-wide MHC-I genetic diversity and a small number of MHC-I alleles compared to outbred mainland populations. However, each individual Raso lark had a rather large number of MHC-I alleles and these alleles were highly divergent. The explanation for the discrepancy between population and within-individual estimates is that the within-locus genetic diversity is low, as shown by a high degree of homozygosity, whereas the gene copy number is high. Overall the immune gene diversity in the genome of Raso larks is surprisingly high, which is promising for this threatened species.

### 4.1 Inheritance and genomic architecture of MHC-I loci

The three fixed or near fixed alleles (A, B, C) most likely represent three core MHC-I loci, not only due to their commonness but also because of their relatively high divergence (average amino acid distance 0.364 ± 0.068; Figure 1a; Figure S4,). The co-segregating block of five alleles (*EFGHK*) occurring in 52% of the individuals contributed an increased MHC-I diversity by 50% compared to individuals without this co-segregating block. The co-segregating *EFGHK* block had high within-block divergence (average amino acid distance 0.356 ± 0.039), though as expected the individual alleles within the co-segregating block had low divergence when compared pairwise with the alleles A, B and C: amino acid distance A–G 0 (1 synonymous substitution; and A–P 0.012 with 1 non-synonymous substitution), B–F (and B–S) 0.012 (1 non-synonymous substitution), and C–K 0.037 (three non-synonymous and two synonymous substitutions; Figure S4). We therefore find it likely that the three loci A, B and C have been duplicated and that the corresponding alleles found in the co-segregating *EFGHK* block are the alleles G, F and K (Figure 1a). The two rarer co-segregating blocks *PQU* and *ST* (Figure 1b-c), as well as several single alleles, may also have resulted from duplications (Figure 1a; Table S3).

For the (near) fixed *ABC* alleles and the frequent co-segregating allele block *EFGHK*, the most likely evolutionary scenario is that the A, B and C alleles correspond to loci that have been duplicated and formed the G, F and K loci (Figure 1a; Table S3, Figure S4) – after which further gene duplications have occurred resulting in the E and H loci within the co-segregating block – and that there are two main haplotypes in the population: *ABC* (three loci) and *ABCEFGHK* (eight loci). This is concordant with the observed bimodal ratios of average allele read depth for *EFGHK* over *ABC* alleles in individuals with *EFGHK* alleles, corresponding to two genotypes: heterozygous for *ABCEFGHK*|*ABC* (low ratio, as *EFGHK* occurs with one copy) and homozygous for *ABCEFGHK* (doubled ratio, as *EFGHK* occurs with two copies; Figure 2a). In a linear model, genotype and number of reads for *ABC* alleles explains 93% of the variation in number of reads for *EFGHK* alleles (F_2,56_ = 367, p < 0.001). Further, the observed genotype frequencies are concordant with Hardy-Weinberg equilibrium (58 homozygous for *ABC*, 51 heterozygous for *ABCEFGHK*|*ABC*, 13 homozygous for *ABCEFGHK*; χ^2^ = 0.13, p = 0.72). The alternative interpretation – that the two main haplotypes would be *ABC* and *EFGHK* (where the alleles G, F, and K could be alternative alleles of the A, B, and C loci) – would strongly violate Hardy-Weinberg equilibrium (χ^2^ = 15.42, p < 0.001). This would either imply that we had failed to detect eight *EFGHK* homozygotes or that strong selection led to this deviation from expected genotype frequencies, but neither explanation is consistent with a bimodal distribution of ratios of read depth for *EFGHK* over *ABC* in individuals with *EFGHK* alleles.

Several alleles or co-segregating blocks appear associated with the *ABC* haplotype, whereas allele O co-segregates completely with the *ABCEFGHK* haplotype, save for two occurrences with *ABC* homozygotes (Table S5), which may represent a post-bottleneck recombination event. Consequentially, *ABCEFGHK*|*ABC* heterozygotes have the highest number of alleles and MHC-I diversity (Figure 2b).

The maximum number of MHC-I alleles was 16 (three individuals), which in one individual included the haplotype *ABCEGFHK* (eight loci), the co-segregating block *PQU* (three loci), and five additional alleles (J, N, O, R, V). Given the patterns of co-segregation, this means that the Raso lark genome contains at least 14 (8+3+6/2) different functional MHC-I loci.

### 4.2 Maintenance of high within-individual MHC-I diversity in a severely bottlenecked population

We identified 22 MHC-I alleles in the Raso lark, many of which were present at high frequency in the sampled population, and often co-occurred within individuals at high levels, corresponding to at least 14 MHC-I loci. Further, based on our evaluation of multiple primer pairs (see Material and Methods), we are certain that other primer pairs recovered one more allele with high frequency and coverage (additional allele 1; not clustering closely with any of the present alleles), as well as seven other functional alleles at low to very low coverage (alleles 2–8; none of which clustering closely with any of the present alleles, but three of which clustering tightly together), see Figure S2. It is thus possible that there are as many as 31 MHC-I alleles in the Raso lark. Including such low-coverage alleles would also increase the minimum number of MHC-I loci to 15 (additional alleles 1 and 2 detected in one of the individuals with 16 main alleles).

Minias, Pikus, Whittingham, Dunn, and Niimura (2019) recently reviewed copy number variation of MHC loci across the avian tree, and among 35 characterized species, only in four cases were more loci established than in the Raso lark. Even when restricting that dataset to species in which >100 individual have been genotyped, a higher number of loci had only been established in two: (1) Based on genotyping of 857 individuals, the great tit *Parus major* displayed no less than 862 functional alleles, and Sepil, Moghadam, Huchard, and Sheldon (2012) could establish at least 16 functional MHC-I loci. (2) The sedge warbler *Acrocephalus schoenobaenus* holds the record so far, with an astounding >3,500 MHC class I alleles detected in 863 individuals, with a within-individual diversity of 65 alleles and thus at least 33 loci (Biedrzycka et al., 2017). In addition, (3) in a recent study Roved, Hansson, Stervander, Hasselquist, and Westerdahl (in prep) not only characterized a high allelic diversity of 390 alleles in 548 great reed warblers, but also inferred haplotypes – the largest of which comprised 17 loci. In contrast, in the widespread blue tit 50 alleles were detected in 918 individuals, with a maximum within-individual diversity of 19 alleles, indicating a minimum of 10 loci (Wutzler, Foerster, & Kempenaers, 2012).

Further, comparing 12 (O’Connor et al., 2016) and 32 (O’Connor, Cornwallis, Hasselquist, Nilsson, & Westerdahl, 2018) species, but sequencing only three and four individuals of each, O’Connor et al. could demonstrate a strong phylogenetic signal in individual allele count, with large differences between different passerine lineages ranging from single digits to around 40. While resident sub-Saharan African species generally had higher MHC-I diversity than their resident Palaearctic or trans-continental migratory relatives (O’Connor et al., 2018), there is no evolution towards a global diversity optimum; instead the evolution of diversity is concordant with fluctuating selection or genetic drift (O’Connor et al., 2016). While it is remarkable that the severely bottlenecked Raso lark has preserved a number of MHC-I loci that equals or exceeds that of several other widespread species of much larger contemporary populations, the Raso lark is part of the superfamily Sylvioidea (including, e.g., many warbler families, swallows, white-eyes, and larks), which displays the highest MHC-I allele counts among birds (Minias, Pikus, Whittingham, et al., 2019). Regrettably, we know of no study of MHC-I diversity in the widespread congeners Eurasian skylark *A. arvensis* or Oriental skylark *A. gulgula*, or for that matter any other lark (family Alaudidae). Although the number of MHC-I loci is high in the Raso lark, the total number of alleles is low, and the degree of allele sharing is high. We compared the frequency in a population for the Raso lark alleles with alleles of three outbred species: Eurasian siskin *Spinus spinus* (Drews & Westerdahl, 2019), house sparrow, and great reed warbler *Acrocephalus arundinaceus* (Roved, Hansson, Tarka, Hasselquist, & Westerdahl, 2018). These three species all have rather few high-frequency alleles and long tails of many low-frequency alleles, whereas the pattern is the opposite for the Raso lark (Figure 4). Interestingly, an intermediate distribution was found by Gonzalez-Quevedo, Phillips, Spurgin, and Richardson (2015) in another island endemic, the Berthelot’s pipit *Anthus berthelotii*. However, while this species has passed through an initial bottleneck during colonization from the mainland, and subsequent ones during between-island colonisations around 75 thousand years ago (Spurgin et al., 2011), it is now widespread with a large population over 13 islands in three archipelagos.

**Figure 4.**
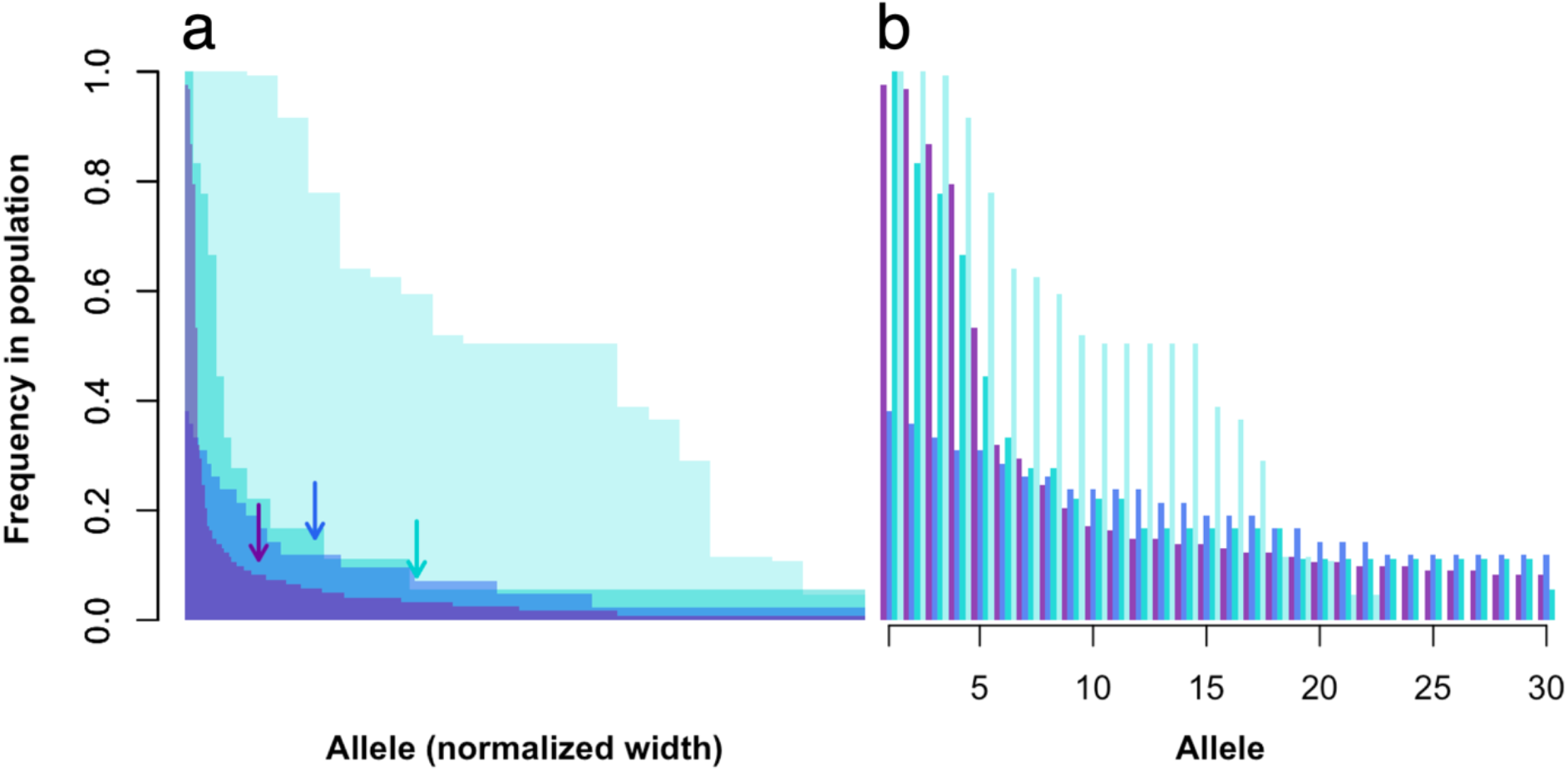
Allele frequency distribution in the Raso lark *Alauda razae* (light turquoise; 22 alleles in 122 individuals; this study), compared to populations of Eurasian siskin Spinus spinus (dark turquoise; 88 alleles in 18 individuals; Drews and Westerdahl (2019)), house sparrow Passer domesticus (blue; 157 alleles in 42 individuals; unpublished data), and great reed warbler Acrocephalus arundinaceus (purple; 271 alleles in 122 individuals—the same number as Raso larks—randomly selected from Roved et al. (2018)). Alleles are ordered along the x-axis following decreasing frequency in the populations. (**a**) The width of individual allele bars has been normalized based on total number of alleles. Thus, while the number of alleles differ between the species, the distributions are directly comparable. (**b**) Frequency of the 30 most frequent alleles displayed, grouped by species. Arrows in (**a**) indicates the position of the 30^th^ allele, except for the Raso lark that only has 22 alleles.

In comparison with other bottlenecked populations, the Raso lark displays high diversity. The only passerine where there is data to draw comparison is the Seychelles warbler (also within the superfamily Sylvioidea), which went through a population bottleneck of less than 30 individuals in the 1960s but has since recovered to a current estimate of 3,000 individuals. This species only has 10 alleles over at least four loci (Richardson & Westerdahl, 2003). Among bottlenecked non-passerines, the white-tailed eagle *Haliaeetus albicilla* (1,000 pairs in the 1970s) has only 10 MHC-I alleles over at least three loci (Minias, Pikus, & Anderwald, 2019), the vulnerable Chinese egret *Egretta eulophotes* (declining population of 2,600–3,400 individuals) has 14 alleles but shows little polymorphism in presumably only two loci (Chiang et al., 2017), and 16 alleles over 6 loci were found in the Endangered red-crowned crane *Grus japonica* (declining population of 1,830 individuals) (Akiyama et al., 2017). However, it is important to note that, in general, non-passerine birds have fewer MHC loci (Minias, Pikus, Whittingham, et al., 2019). For example, both in falcons *Falco* spp. (Gangoso et al., 2012) and prairie-chickens *Tympanuchus* spp. (Bateson, Whittingham, Johnson, & Dunn, 2015; Minias et al., 2016), species listed either as Threatened or as Least Concern all displayed a single MHC-I locus. Furthermore, one should acknowledge that it is difficult to design primers that amplify all potential alleles in a hypervariable part of the genome, as we could demonstrate with our evaluation of newly designed primer pairs, and the number of alleles reported for various species may therefore be underestimated.

Recent results from Dierickx, Sin, et al. (2019) provide a possible explanation for the relatively high level of MHC-I diversity in the Raso lark. They found that, while the Raso lark did indeed have reduced genetic diversity compared to its widespread, continental, closest relatives, this difference was smaller than expected. Their results show that this can be largely explained by the recency of a population contraction from a much larger ancestral size, an explanation that could also be valid for MHC-I diversity.

However, despite the considerable genetic diversity in Raso larks relative to their population size, Dierickx, Sin, et al. (2019) point out that the species is still in a precarious situation: continued existence at this small population size will unavoidably increase their genetic risks, including inbreeding.

### 4.3 Effects of MHC-I diversity on mate choice and fitness

Based on the above, we hypothesised that female Raso larks would choose males with MHC genotypes complementary to theirs, in order to maximise the genetic diversity of their offspring. However, we found no evidence for this phenomenon. One explanation for the absence of effect of MHC complementarity on mate choice is that, given that the levels of genetic diversity are still relatively high in the population (Dierickx, Sin, et al., 2019), and that the population contraction around 550 years ago, coinciding with human settlement, was recent from an evolutionary point of view (Dierickx, Sin, et al., 2019), maybe there has been no strong selection yet to maximise diversity through mate choice; maybe not enough time has passed yet for that behaviour to evolve.

It is also possible that other mating criteria override the importance of MHC in mate choice. For example, males occupy and defend territories, and these could play a role in the females’ choice (e.g. Alatalo, Lundberg, & Glynn, 1986; Bensch & Hasselquist, 1992; Buchanan & Catchpole, 1997). Indeed, in mating systems where there are many indirect benefits to mating, females might care less about the genetic make-up of the male (Zelano & Edwards, 2002). Richardson and Komdeur (2005) did not find any influence of MHC on social partner choice in female Seychelles warblers (Richardson, Komdeur, Burke, & von Schantz, 2005). The social partner probably provides enough indirect benefits (e.g. nest building, feeding of the chicks) to the female for her not to take MHC diversity into account when choosing him. However, Richardson and Komdeur (2005) found that, when the social partner had a low level of MHC diversity, the female was more likely to engage in extra-pair copulations, with a male that had a higher level of MHC diversity than the partner (Richardson et al., 2005). We do not know enough about the details of the Raso lark’s socially monogamous mating system to be able to hypothesize whether a similar phenomenon might be at play in our study system. Our results are based on social pairings. It is possible that female Raso larks engage in extra-pair matings, and indeed the frequency of extrapair offspring is 20 percent in a close relative of the Raso lark, the Eurasian skylark (Hutchinson & Griffith, 2008).

A final functional explanation for the absence of MHC-based mate choice might be due to the fact that larks in arid environments generally have very few parasites (Horrocks et al., 2012). This would reduce the importance of the immune system for survival and reproductive fitness and thereby reduce selection for MHC-based mate choice. Our analyses did not find any effect of MHC-I diversity on survivorship. That this is the case though does then beg the question of why the Raso lark has such high levels of MHC-I diversity, if not to fight a wide array of diseases. The high diversity may reflect past selection, as the Raso lark probably diverged from the ancestor of two currently widespread continental species, the skylark and the oriental skylark *A. gulgula*, about six million years ago (Alström et al., 2013), and nothing is known about MHC-I diversity in the congeneric species. Further, based on our results, it seems as though while within-locus diversity may be low (i.e. there seems to be a single allele for many loci), between-locus diversity is high, and overall functional MHC-I diversity has been maintained through population bottlenecks at a high level by the co-segregation of large blocks of linked loci (e.g. the haplotype *ABCEFGHK*).

### 4.4 Conclusions

This study shows that it is possible for a population to maintain relatively high levels of MHC diversity even in the face of extremely severe bottlenecks, facilitated by the co-segregation of large numbers of linked divergent loci (gene copies), despite a low number of total alleles. This remaining diversity will likely serve the Raso lark well when adapting to future environmental change (cf. de Villemereuil et al., 2019), especially in the context of the ongoing reintroduction of the species to nearby Santa Luzia Island and of Cape Verde’s vulnerability in the face of climate change.

## Supporting information

Supporting Information 3 (allele sequence alignment, fasta format)

## Acknowledgements

Anna Drews offered invaluable help with amplicon library preparations. Thanks to the field assistants on Raso: Mark Bolton, Ewan Campbell, Simon Davies, Mike Finnie, Tom Flower, Lee Gregory, Sabine Hille, Mark Mainwaring, Jason Moss and Justin Welbergen. This work has been generously supported by the Sir Peter Scott Studentship and the Rouse Ball Eddington Fund of Trinity College, Cambridge (to EGD), the VOCATIO Award (to EGD), the European Research Council (ERC) under the European Union’s Horizon 2020 research and innovation programme with grants 679799 (to HW), and Julian Francis, RSPB, CEPF and BirdLife International’s Preventing Extinctions Initiative.

## Author contributions

EGD devised the research questions and hypotheses, EGD, MS and HW designed the study, MLB and EGD collected the Raso lark samples, MS did the laboratory work, bioinformatics, and MHC characterization analyses, JT carried out the mate choice and survivorship statistics, HW provided laboratory resources, and all authors participated in the writing of the paper.

## Data accessibility

Theraw sequence data for 136 samples from two libraries has been submitted to Zenodo, accessible at https://doi.org/10.5281/zenodo.3630877. The 22 alleles A–V and additional alleles 1–2 have been deposited at GenBank with accession numbers MT010367–MT010390.

## SUPPORTING INFORMATION

### Supporting Information 1 (PDF): Supplementary tables and figures

**Table S1:** Sequence, melting temperature, and other details for primers. **Table S2:** PCR conditions and amplification success in four species for 13 primer pairs evaluated in this study. **Table S3:** Properties, such as read counts and presumed parental allele, for alleles A–V and excluded additional alleles 1 and 2. **Table S4:**Genotyped Male Raso larks present in the population in each year of the study. **Table S5:**Association of alleles and co-segregating blocks to *ABCEFGHK*|*ABC* genotypes. **Table S6:** Randomisation testing of (dis-)assortative mating based on allelic variation in MHC-I; corresponds to Table 1, but excludes *RV* alleles. **Figure S1:** Evaluation of amplification success by electrophoresis in four species for 13 primer pairs evaluated in this study. **Figure S2:** Unrooted MHC-I amino acid allele tree, including the alleles called and used for analyses (A–V) and additional alleles that are certainly (alleles 1 & 2) or possibly (alleles 3–9) functional, but yielded too few reads with our primer pairs to enable confident allele calls. **Figure S3:** No relationship between coverage (number of reads) and number of alleles called among the 114 individuals included in the analyses. **Figure S4:** Amino acid and nucleotide distance matrix for the MHC-I exon 3 alleles A–V. **Figure S5:** MHC-I diversity over time, based on inferred birth year. **Figure S6:** Survival probability from time of ringing for nestlings, and adult birds with and without claw damage. **Figure S7:** Randomisation testing of (dis-)assortative mating in Raso larks based on allelic variation in MHC-I; corresponds to Figure 2, but excludes *RV* alleles.

### Supporting Information 2 (XLS)

Data file used for analyses, containing information for 114 individuals on genotype, MHC-I diversity, sex, mate, years observed, years breeding, assumed birth year etc.

### Supporting Information 3 (FA)

Fasta file comprising the 24 MHC class I alleles (A–V, additional alleles 1–2).

