## Supporting Information 1 (Tables S1-S6, Figures S1-S7) for "High MHC gene copy number maintains diversity despite homozygosity in a Critically Endangered single-island endemic bird, but no evidence of MHC-based mate choice"

### Table of Contents:


| | |
|---|---|
| **Table S1:** Sequence, melting temperature, and other details for primers. | **Page 2** |
| **Table S2:** PCR conditions and amplification success in four species for 13 primer pairs evaluated in this study. | **Page 3** |
| **Table S3:** Properties, such as read counts and presumed parental allele, for alleles A–V and excluded additional alleles 1 and 2. | **Page 4** |
| **Table S4:** Evaluation of primer pair combinations, and their PCR conditions. | **Page 5** |
| **Table S5:** Association of alleles and co-segregating blocks to the ABC\|ABCEFGHK genotypes | **Page 6** |
| **Table S6:** Randomisation testing of (dis-)assortative mating based on allelic variation in MHC-I; corresponds to Table 1, but excludes *RV* alleles. | **Page 6** |
| **Figure S1:** Evaluation of amplification success by electrophoresis in four species for 13 primer pairs evaluated in this study. | **Page 7** |
| **Figure S2:** Unrooted MHC-I amino acid allele tree, including the alleles called and used for analyses (A–V) and additional alleles that are certainly (alleles 1 & 2) or possibly (alleles 3–9) functional, but yielded too few reads with our primer pairs to enable confident allele calls. | **Page 8** |
| **Figure S3:** No relationship between coverage (number of reads) and number of alleles called among the 114 individuals included in the analyses. | **Page 9** |
| **Figure S4:** Amino acid and nucleotide distance matrix for the MHC-I exon 3 alleles A–V. | **Page 10** |
| **Figure S5:** MHC-I diversity over time, based on inferred birth year. | **Page 11** |
| **Figure S6:** Survival probability from time of ringing for nestlings, and adult birds with and without claw damage. | **Page 11** |
| **Figure S7:** Randomisation testing of (dis-)assortative mating in Raso larks based on allelic variation in MHC-I; corresponds to Figure 3, but excludes *RV* alleles. | **Page 12** |

**Table S1** MHC class I exon 3 primers evaluated for this study. For the primers developed for this project, the naming indicates exon (3), direction (F/R), 5' starting position in relation to exon boundary (us = upstream, ds = downstream; us17 meaning 17 nucleotides upstream of the outer exon nucleotide), and 3' end position in relation to exon boundary (ex = exon; ex6 meaning the 6th nucleotide from the start/end of the exon). Melting temperature ($T_M$).

| **Primer name** | **Direction** | **Primer sequence 5'–3'** | **$T_M$ (°C)** | **Reference** |
|---|---|---|---|---|
| 3F_us17-ex10 | forward | GGGTACAATCCCCCCAGGTCTYCACAC | 68.2–69.7 | this study |
| 3F_us30-us4 | forward | CATGGGTCTCTGTGGGTACAATCCCC | 66.4 | this study |
| 3F_us35-us8 | forward | AATTCCATGGGTCTCTGTGGGTACAAT | 64.1 | this study |
| HNalla | forward | TCCCCACAGGTCTCCACAC | 60.8 | Westerdahl et al. (2004) |
| 3R_ds16-ex10 | reverse | AATTCCCACCCACCTTTGCRCTCCAG | 67.2–69.4 | this study |
| 3R_ds20-ex6 | reverse | CACCTTTGCRCTCCAGCTCCTTCTGC | 67.8–69.9 | this study |
| 3R_ds24-ex2 | reverse | AAATCCCAAATTCCCACCCACCTTTG | 64.1 | this study |
| 3R_ds34-ds7 | reverse | CTCCATTCCCAAATCCCAAATTCCCAC | 64.3 | this study |
| MHC1_HN46 | reverse | ATCCCAAATTCCCACCCACCTT | 61.9 | Westerdahl et al. (2004) |

**Table S2** Genotyped Male Raso Larks present in the population in each year of the study.

| Year | Number of individuals |
|---|---|
| 2004 | 3 |
| 2005 | 6 |
| 2006 | 11 |
| 2007 | 13 |
| 2008 | 16 |
| 2009 | 19 |
| 2010 | 28 |
| 2011 | 30 |
| 2012 | 33 |
| 2013 | 31 |
| 2014 | 24 |
| 2015 | 23 |
| 2016 | 18 |
| 2017 | 1 |

**Table S3** Allele properties. Properties and read counts of the 22 MHC-I exon 3 alleles A–V recovered by amplicon sequencing of Raso Lark *Alauda razae*. "Parental alleles" are defined as alleles of higher read count that differ ≤2 nucleotides (distance specified within parentheses). Allele depth corresponds to the total count of read pairs across samples; N samples are the number of samples, out of the original 130 samples (of 122 individuals, including eight technical replicates) with >5,000 reads, in which the allele was recovered; Min count represent the minimum number of read pairs for that allele recovered in a single individual; Mean count represents the average count of read pairs in individuals with that allele. The colours indicate co-inherited blocks of alleles; cf. Figure 1, Figure S4, Figure S2. *Denotes two alleles of much lower per-individual coverage; see the main text. Two additional functional alleles (alleles 1 and 2) had too low read depth for reliable allele calling with the main primers used. All allele sequences are available on GenBank with accession numbers MT010367–MT010390 and in Supporting Information 3 (fasta alignment).

| Allele name | Parental allele | Formal allele name | Allele depth | N samples | Min count | Mean count |
|---|---|---|---|---|---|---|
| A | | Alra-UA*01 | 730,748 | 130 | 1,114 | 5,621 |
| B | | Alra-UA*02 | 585,095 | 129 | 1,215 | 4,536 |
| C | | Alra-UA*03 | 376,909 | 130 | 659 | 2,899 |
| D | | Alra-UA*04 | 222,367 | 81 | 485 | 2,745 |
| E | | Alra-UA*05 | 168,200 | 66 | 1,082 | 2,548 |
| F | B (1) | Alra-UA*06 | 141,592 | 66 | 957 | 2,145 |
| G | A (1) | Alra-UA*07 | 141,934 | 66 | 949 | 2,151 |
| H | | Alra-UA*08 | 128,678 | 66 | 865 | 1,949 |
| K | | Alra-UA*09 | 87,454 | 66 | 596 | 1,325 |
| I | | Alra-UA*10 | 121,246 | 48 | 581 | 2,578 |
| J | D (1) | Alra-UA*11 | 99,123 | 51 | 560 | 1,944 |
| L | | Alra-UA*12 | 65,866 | 38 | 448 | 1,733 |
| M | | Alra-UA*13 | 59,629 | 83 | 149 | 718 |
| N | M (1) | Alra-UA*14 | 44,890 | 78 | 99 | 576 |
| O | | Alra-UA*15 | 43,033 | 68 | 142 | 633 |
| P | A (1); G (2) | Alra-UA*16 | 36,383 | 15 | 1,234 | 2,426 |
| Q | H (1) | Alra-UA*17 | 32,775 | 14 | 1,256 | 2,341 |
| U | | Alra-UA*18 | 19,672 | 15 | 647 | 1,311 |
| R* | | Alra-UA*19 | 32,627 | 117 | 47 | 281 |
| S | B (1); F (2) | Alra-UA*20 | 22,397 | 6 | 1,804 | 3,733 |
| T | I (1) | Alra-UA*21 | 20,446 | 6 | 1,794 | 3,408 |
| V* | | Alra-UA*01 | 14,345 | 99 | 23 | 142 |
| **1** | | – | 2,641 | 30 | 15 | 75 |
| **2** | | – | 666 | 6 | 5 | 27 |

**Table S4** Evaluation of primer pair combinations, and their PCR conditions. Primer pair ID corresponds to the numbers in Figure S1. Annealing temperature ($T_A$). Successful specific (+), unspecific (m) or no (–) amplification was scored in great reed warbler *Acrocephalus arundinaceus* (Aar), Raso lark *Alauda razae* (Ara), blue tit *Cyanistes caeruleus* (Ccae), and zebra finch *Taeniopygia guttata* (Tgu), see Additional file 5. For primer sequences, see Table S1.

| | | | $T_A$ (°C) | | Amplification success | | | |
|---|---|---|---|---|---|---|---|---|
| **Fwd primer** | **Rev primer** | **Primer pair ID** | Qiagen | Phusion | Aar | Ara | Ccae | Tgu |
| 3F_us17-ex10 | 3R_ds24-ex2 | 1 | 62.6 | | + | + | + | – |
| 3F_us17-ex10 | 3R_ds34-ds7 | 2 | 62.6 | | m | – | + | – |
| 3F_us30-us4 | 3R_ds24-ex2 | 3 | 62.6 | 72 | + | + | + | – |
| 3F_us30-us4 | 3R_ds34-ds7 | 4 | 62.6 | | m | – | + | – |
| 3F_us35-us8 | 3R_ds16-ex10 | 5 | 62.6 | | + | + | + | + |
| 3F_us35-us8 | 3R_ds20-ex6 | 6 | 62.6 | 72 | + | + | + | + |
| 3F_us35-us8 | 3R_ds24-ex2 | 7 | 62.6 | 72 | + | + | + | – |
| 3F_us35-us8 | 3R_ds34-ds7 | 8 | 62.6 | | m | – | m | – |
| 3F_us30-us4 | 3R_ds16-ex10 | 9 | 64.6 | | + | + | + | + |
| 3F_us30-us4 | 3R_ds20-ex6 | 10 | 64.6 | | + | + | + | + |
| 3F_us17-ex10 | 3R_ds16-ex10 | 11 | 65.7 | | + | + | + | + |
| 3F_us17-ex10 | 3R_ds20-ex6 | 12 | 65.7 | | + | + | + | + |
| MHC1_HNalla | MHC1_HN46 | 26 | 59.3 | 72 | + | + | + | na |

**Table S5** Association of alleles and co-segregating blocks to the ABC/ABCEFGHK genotypes. Alleles or co-segregating blocks that seem to associate with the ABC haplotype (no observation in ABCEFGHK homozygotes) are highlighted blue, whereas one allele, O, with association to the ABCEFGHK haplotype (observed in all individuals ABCEFGHK and two ABC homozygotes) is highlighted pink. Significance (Sign.) levels corresponding to α = 0.05 (*), 0.01 (**), and 0.001 (***) after Bonferroni correction for multiple $\chi^2$ tests.

| | Genotype ABC \| ABCEFGHK | | | |
|---|---|---|---|---|
| **Allele(s)** | hom ABCEFGHK | het | hom ABC | **Sign.** |
| ABC / ABCEFGHK | 13 | 51 | 58 | |
| D | 3 | 28 | 45 | |
| I | 0 | 17 | 28 | |
| J | 9 | 34 | 8 | *** |
| L | 8 | 18 | 9 | |
| M | 0 | 30 | 47 | * |
| N | 13 | 43 | 18 | ** |
| O | 13 | 51 | 2 | *** |
| PQU | 0 | 1 | 13 | * |
| ST | 0 | 1 | 4 | |
| R | 11 | 48 | 50 | |
| V | 0 | 36 | 58 | ** |

**Table S6** Randomisation testing of (dis-)assortative mating based on allelic variation in MHC-I, excluding *RV* alleles. The empirical means calculated from observed pairings were compared to randomly assigned pairings generated from 9,999 permutations; significant departures from random mating would fall outside the 95% confidence intervals. α and β refer to the whether the co-segregating block of five alleles were treated as five separate alleles (α), or a single allelic block (β). *p* is the ranked position of the observed mean in the distribution.

| Variable | Empirical mean | 2.5% CI | 97.5% CI | *p* |
|---|---|---|---|---|
| **Excluding *RV* alleles** | | | | |
| Allele sharing α | 0.634 | 0.591 | 0.699 | 0.357 |
| Allee sharing β | 0.565 | 0.538 | 0.614 | 0.304 |
| Mean amino acid distance | 13.85 | 13.61 | 13.92 | 0.852 |

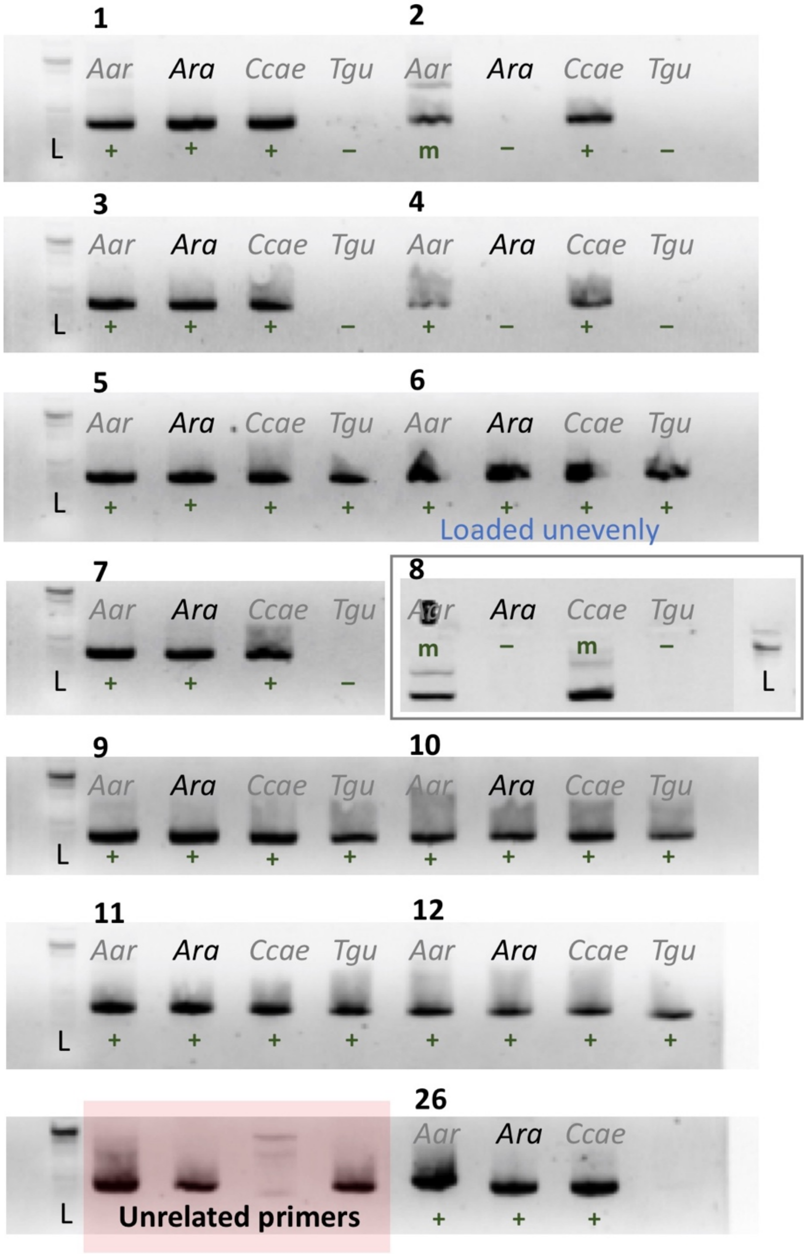


**Figure S1** Evaluation of PCR success by electrophoresis on 2 % agarose gels with a Gibco 1 Kb ladder (L) for different primer combinations (numbers; see Table S2). The samples used for evaluation were great reed warbler *Acrocephalus arundinaceus* (Aar), Raso lark *Alauda razae* (Ara), blue tit *Cyanistes caeruleus* (Ccae), and zebra finch *Taeniopygia guttata* (Tgu). Successful specific (+), unspecific (m) or no (–) amplification is scored under the DNA bands. Note that primer combination 8 was evaluated on a separate gel, which was run for a longer time.

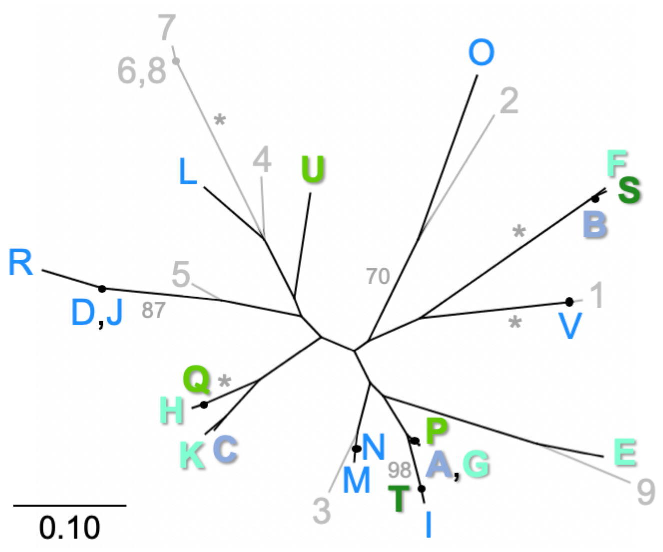


**Figure S2** Unrooted MHC-I amino acid allele tree, including the alleles called and used for analyses (A–V; black branches) and additional alleles that are certainly (alleles 1 & 2) or possibly (alleles 3–9) functional (all with grey branches and names), but yielded too few reads with our primer pairs to enable confident allele calls. The tree is computed with the LG substitution model, and corresponds to Figure 1a (which does not include additional alleles 1–9). Allele names are placed at tips or internal terminal nodes (marked with black circles). Three pairs (A/G, D/J, and 6/8) do not differ by any non-synonymous mutations (see Additional file 1: Figure S5). Fixed or almost fixed alleles, as well as alleles in co-segregating clusters, are highlighted in bold and coloured as in Figure 1, Table S3, and Figure S5 (Additional file 1). Bootstrap values >70%, based on 1,000 replicates, are written with grey font (at the branch upstream of supported node), with 100% indicated by an asterisk.

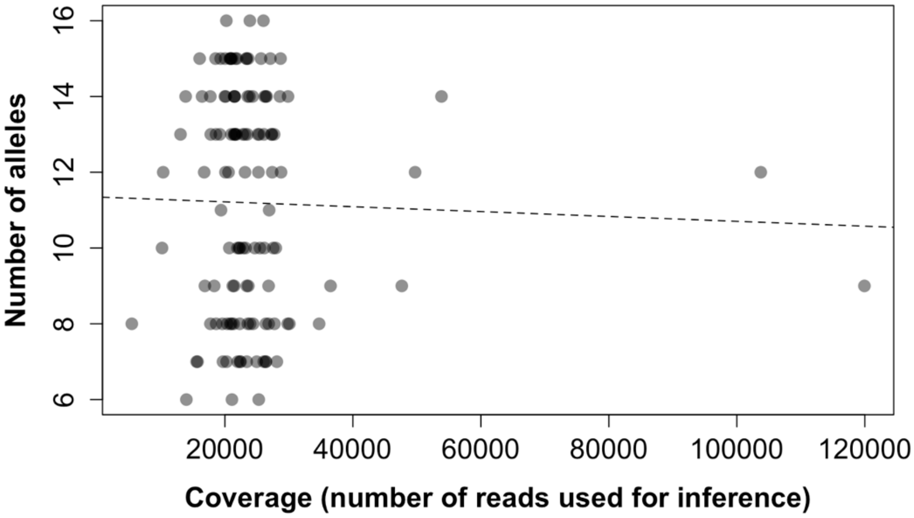


**Figure S3** Among the 122 samples retained in the analysis, there was no effect of effective coverage on the number of alleles inferred. The flat dashed line corresponds to a non-significant linear regression.

| | A | B | C | D | E | F | G | H | K | I | J | L | M | N | O | P | Q | U | R | S | T | V |
|---|---|---|---|---|---|---|---|---|---|---|---|---|---|---|---|---|---|---|---|---|---|---|
| A | | **35** | **27** | 33 | 26 | 36 | 1 | 28 | 28 | 8 | 32 | 28 | 10 | 11 | 31 | 1 | 27 | 23 | 34 | 36 | 7 | 30 |
| B | **21** | | **40** | 37 | 33 | 1 | 36 | 37 | 35 | 37 | 38 | 38 | 29 | 30 | 40 | 34 | 36 | 36 | 41 | 1 | 36 | 36 |
| C | **15** | **23** | | 27 | 35 | 41 | 28 | 17 | 5 | 28 | 26 | 25 | 28 | 27 | 37 | 26 | 16 | 24 | 25 | 41 | 29 | 38 |
| D | 17 | 23 | 12 | | 41 | 38 | 34 | 32 | 24 | 33 | 1 | 34 | 29 | 28 | 32 | 32 | 31 | 31 | 6 | 38 | 34 | 40 |
| E | 15 | 19 | 20 | 23 | | 34 | 27 | 35 | 34 | 30 | 40 | 37 | 26 | 27 | 39 | 25 | 34 | 40 | 42 | 32 | 29 | 33 |
| F | 21 | 1 | 23 | 24 | 19 | | 37 | 38 | 36 | 38 | 39 | 39 | 30 | 31 | 39 | 35 | 37 | 37 | 42 | 2 | 37 | 37 |
| G | 0 | 21 | 15 | 17 | 15 | 21 | | 29 | 29 | 9 | 33 | 29 | 11 | 12 | 32 | 2 | 28 | 24 | 35 | 37 | 8 | 31 |
| H | 18 | 22 | 9 | 17 | 21 | 23 | 18 | | 18 | 27 | 31 | 29 | 29 | 30 | 37 | 27 | 1 | 25 | 30 | 38 | 28 | 41 |
| K | 16 | 20 | 3 | 11 | 19 | 21 | 16 | 10 | | 29 | 23 | 24 | 25 | 24 | 38 | 27 | 17 | 23 | 24 | 36 | 30 | 35 |
| I | 6 | 22 | 15 | 17 | 18 | 22 | 6 | 18 | 16 | | 32 | 27 | 16 | 17 | 36 | 7 | 26 | 28 | 32 | 38 | 1 | 38 |
| J | 17 | 23 | 12 | 0 | 23 | 24 | 17 | 17 | 11 | 17 | | 33 | 30 | 29 | 31 | 31 | 30 | 32 | 5 | 39 | 33 | 39 |
| L | 17 | 21 | 16 | 18 | 21 | 22 | 17 | 19 | 15 | 17 | 18 | | 30 | 29 | 34 | 27 | 28 | 21 | 35 | 39 | 28 | 41 |
| M | 6 | 18 | 16 | 17 | 14 | 18 | 6 | 17 | 15 | 10 | 17 | 17 | | 1 | 33 | 11 | 28 | 21 | 33 | 30 | 15 | 31 |
| N | 7 | 19 | 15 | 16 | 15 | 19 | 7 | 18 | 14 | 11 | 16 | 16 | 1 | | 34 | 12 | 29 | 20 | 32 | 31 | 16 | 30 |
| O | 17 | 26 | 22 | 20 | 23 | 25 | 17 | 22 | 24 | 20 | 20 | 20 | 17 | 18 | | 30 | 36 | 34 | 32 | 41 | 35 | 41 |
| P | 1 | 20 | 14 | 16 | 14 | 20 | 1 | 17 | 15 | 5 | 16 | 16 | 7 | 8 | 16 | | 26 | 24 | 33 | 35 | 6 | 31 |
| Q | 17 | 21 | 8 | 16 | 20 | 22 | 17 | 1 | 9 | 17 | 16 | 18 | 16 | 17 | 21 | 16 | | 24 | 29 | 37 | 27 | 40 |
| U | 13 | 22 | 14 | 18 | 22 | 22 | 13 | 17 | 15 | 17 | 18 | 10 | 13 | 12 | 19 | 14 | 16 | | 33 | 37 | 29 | 32 |
| R | 20 | 25 | 11 | 4 | 24 | 26 | 20 | 17 | 12 | 18 | 4 | 20 | 21 | 20 | 21 | 19 | 16 | 20 | | 40 | 33 | 40 |
| S | 21 | 1 | 23 | 23 | 19 | 2 | 21 | 22 | 20 | 22 | 23 | 21 | 18 | 19 | 26 | 20 | 21 | 22 | 25 | | 37 | 37 |
| T | 5 | 21 | 15 | 17 | 17 | 21 | 5 | 18 | 16 | 1 | 17 | 17 | 9 | 10 | 19 | 4 | 17 | 17 | 18 | 21 | | 37 |
| V | 15 | 20 | 21 | 20 | 19 | 21 | 15 | 22 | 20 | 20 | 20 | 19 | 16 | 15 | 21 | 16 | 21 | 13 | 23 | 20 | 19 | |

**Figure S4** Distance matrix for the 22 MHC-I exon 3 alleles. Above the diagonal are nucleotide distances, below the diagonal amino acid distances. Cell shading highlights more similar alleles (darker). The outlined coloured quadrats indicate co-inherited alleles (see corresponding shading colours in Figure 1, Table S3, and Figure S2) and the bold type indicates fixed or near-fixed alleles.

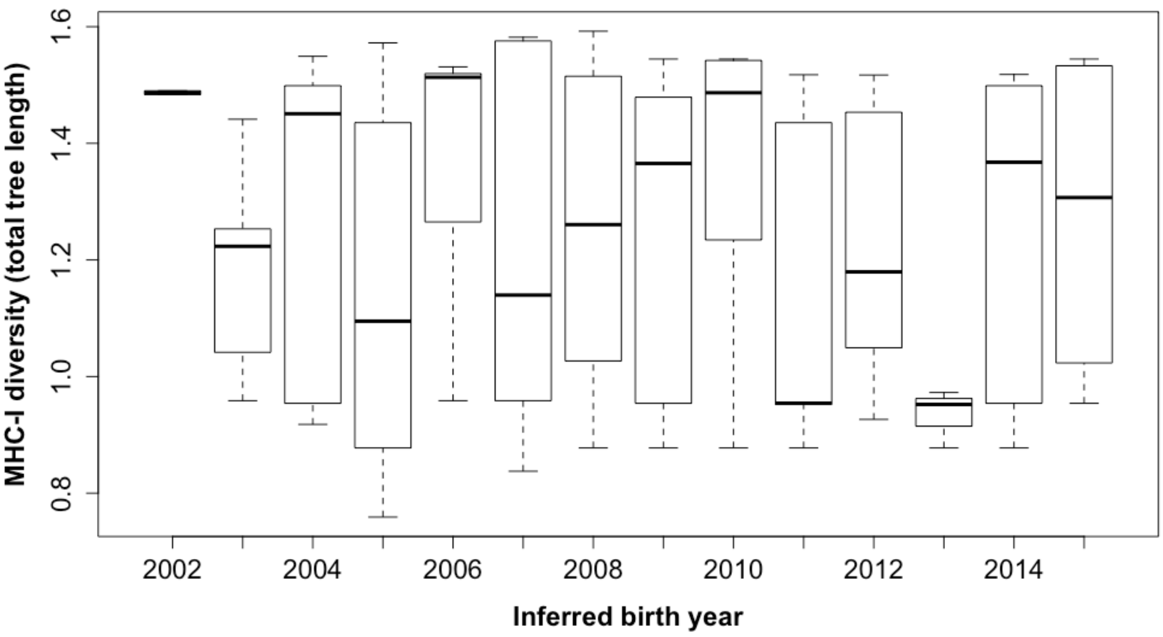


**Figure S5** No trend of MHC-I diversity over time, based on birth year inferred by claw damage (see main text).

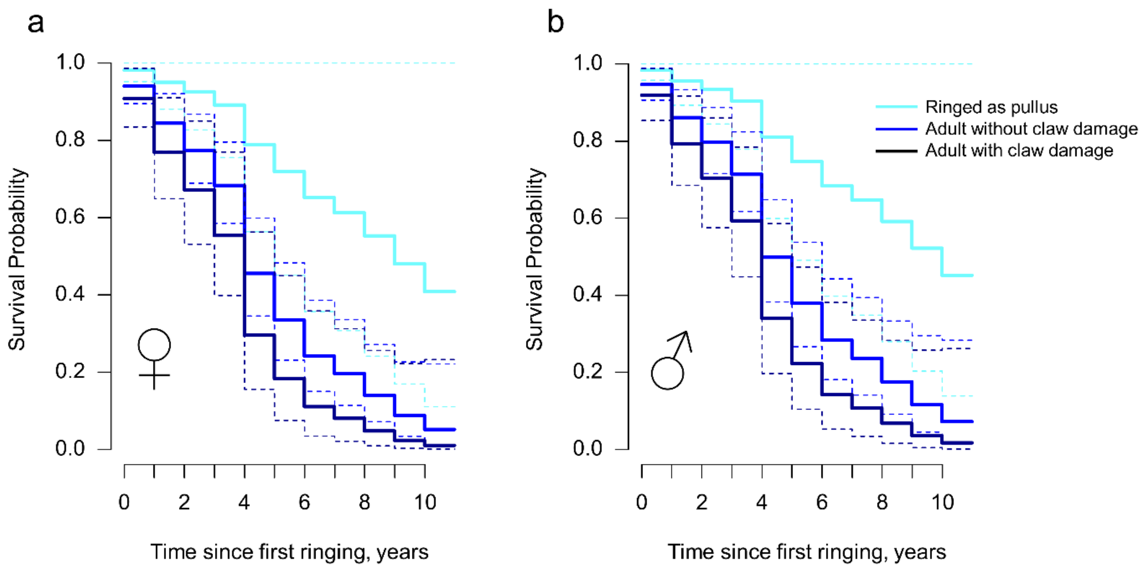


**Figure S6** Survival probability from the time of first ringing for (a) female and (b) male Raso larks according to age categorization. Solid lines represent predicted survival probability from the cox proportional hazards model, with 95% confidence intervals displaying the 95% confidence intervals. For predictions the MHC-1 diversity (the total tree length per genotype), which was non-significant in the model, was set to the population-level mean.

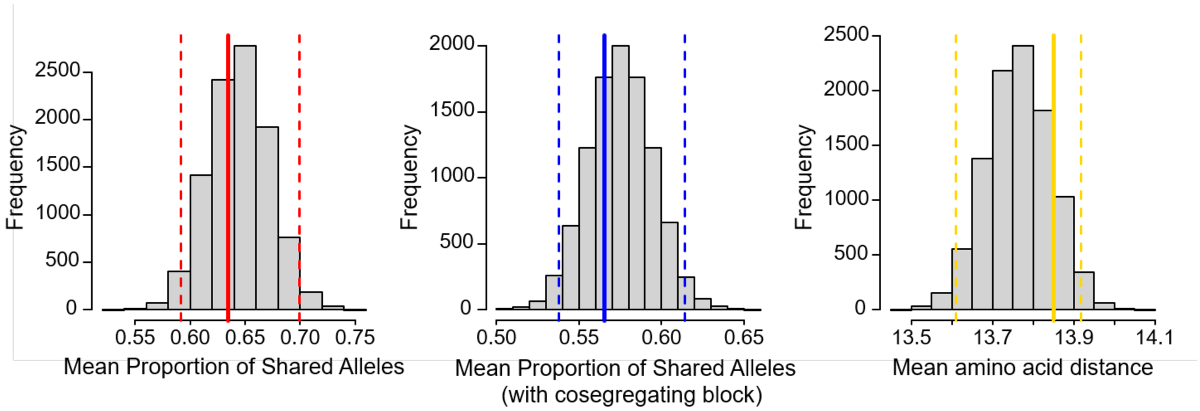


**Figure S7** Randomisation testing of (dis-)assortative mating in Raso larks based on allelic variation in MHC-I. Solid lines show empirical means from observed pairings. Frequency distributions show mean values generated from 9,999 permutations of random pairing (including the empirical data = 10,000). The two-tailed 95% confidence intervals (dashed lines) display cut-offs for significant departures from random mating. n = 46 pairings. This figure excludes the 'RV' alleles; for the corresponding figure including those, see Figure 2.
